# MACS: Multi Domain Adaptation Enables Accurate Connectomics Segmentation

**DOI:** 10.1101/2025.09.30.679580

**Authors:** Abrar Rahman Abir, Anik Saha, Md Shamsuzzoha Bayzid

## Abstract

Connectomics aims to map the brain’s neural wiring by segmenting cellular structures from high-resolution electron microscopy (EM) images. Manual labeling and proofreading remain a major bottleneck for accurate extraction of microstructures. While computational models have advanced automated segmentation, they typically require training from scratch on each dataset, demanding substantial annotated data. Domain adaptation methods address this by transferring knowledge from a labeled source to a less-annotated target. However, existing approaches are limited to adaptation from a single source domain. This overlooks the potential benefits of integrating information from multiple diverse domains, motivating the development of multidomain adaptation. To address this, we propose MACS, the first known multi-domain adaptation framework that combines knowledge from multiple heterogeneous source domains to learn segmentation in the target domain, and employs active learning to efficiently select the most informative target samples for annotation. MACS uses information-theoretic weighting to combine source domains, and introduces a novel and efficient Bayesian Laplace approximation for uncertainty estimation. Our extensive experiments across nine connectomics datasets demonstrate that MACS consistently and substantially outperforms state-of-the-art models, even under limited annotation budgets, with a mean improvement of 5.89% at the lowest annotation budget and 27.72% at the highest annotation budget. In-depth analyses further reveal that MACS offers mechanistic interpretability by quantifying and explicitly upweighting the most transferable source domains for each target.

The preprocessed datasets and the source code of MACS are publicly available at http://github.com/abrarrahmanabir/MACS.

## 1 Introduction

Connectomics is a field aimed at deciphering the wiring diagram of the brain by mapping synaptic joints from high-resolution imaging data. Accurate segmentation of neurons, cell-membranes, and synapses from electron microscopy (EM) images of brain tissue volumes comprises a major part of this workflow. [1] However, extracting such micro-structures from nanometer-scale images requires substantial human efforts in labeling and proofreading, thereby making segmentation a challenging bottleneck of the entire procedure.

The acquisition of connectomics data is rapidly approaching an exabyte scale [2]. However, annotation of such large volumes by human experts is practically infeasible [3]. Nor can we completely rely on automated methods yet due to the prevalence of merge and split errors [4]. The reliance of most connectomics projects on scratch training further exacerbates the issue. These methods not only escalate the annotation costs but also limit the prospect of generalization across datasets.

To mitigate the challenges of scratch-training, transfer learning and domain adaptation have emerged as effective strategies to bridge domain shift. These approaches enable the use of annotated data from a source domain to train models for a target domain, often with minimal or no labeled data. In connectomics, [5] and [6] demonstrates the efficacy of unsupervised domain adaptation in mitochondria and synapse segmentation with predefined source and target domains. In contrast, [7] proposes NeuroADDA, which chooses an optimal source domain from multiple candidates for a given target domain. This approach outperforms scratch training leveraging active learning with a constrained annotation budget in the target domain. However, single-source domain adaptation methods, such as those described, limit the potential for simultaneous alignment of multiple source domains with the target, thereby restricting generalization. Furthermore, traditional domain adaptation frameworks in connectomics typically require that all source domain models share an identical architecture [7]. This restriction severely limits real-world applicability, as publicly available pretrained models are rarely standardized, and optimal performance on diverse datasets often necessitates the use of different model designs.

To address the limitations of scratch-training and single-source domain adaptation, we propose a novel framework **MACS** (***M***ulti Domain ***A***daptation for Accurate ***C*** onnectomics ***S*** egmentation), which is the first attempt at integrating multi-source domain adaptation with active learning to enhance the precision of connectomics image segmentation. Our main contributions are as follows:

1. We propose MACS, a novel multi-domain adaptation framework for connectomic segmentation, capable of leveraging multiple heterogeneous source domains and model architectures.
2. We introduce a novel integration of active learning with analytic Bayesian Laplace-based uncertainty estimation in MACS, enabling both efficient sample selection for annotation. This approach provides scalable uncertainty quantification and significantly reduces computational overhead compared to traditional Monte Carlo dropout, representing a significant advance in uncertainty estimation for large-scale segmentation tasks.
3. We provide extensive empirical validation across nine major connectomics datasets, benchmarking MACS against six representative baselines where MACS consistently and significantly outperforms the baseline models and single domain adaptation across diverse domains and annotation budgets. In particular, MACS delivers significant improvements of 50–60% on datasets such as mouse brain, C. elegans, and Octopus Vertical Lobe SFL Tract.
4. MACS supports the integration of heterogeneous model architectures across multiple source domains. This is achieved by aligning source domains and target via a convex combination of individual source kernels and transferring knowledge to a unified student model through weighted ensemble predictions, thus fully decoupling architectural requirements across domains.
5. We provide mechanistic interpretability by demonstrating strong alignment between learned source weights and independent LEEP-based transferability scores.

## 2. Related Work

A wide range of computational methods have emerged for the efficient and accurate segmentation of EM image volumes in connectomics. The architectures used in these methods include 2D-UNet [8, 3, 9], 3D-UNet [10, 11], Mask-RCNN [12], and FFN [13]. Affinity-based agglomeration techniques improve segmentation accuracy[14, 15]. Using Local Shape Descriptors (LSDs) as an auxiliary prediction task further pushes the accuracy and efficiency of connectomics segmentation [16].

However, the need for expert annotation for new domains often poses a challenge for the aforementioned supervised scratch-training methods. Transfer learning and domain adaptation techniques aim to bridge the domain gap by leveraging pretrained models for segmentation in the target domain with minimal annotated samples. A recent study [17] explores the use of transfer learning for cross-dataset and cross-task generalization by training a 3D ResU-Net for connectomic segmentation. They report accurate mitochondria and ER segmentation with limited ground truth annotations. [6] proposes a two-stream U-Net architecture for unsupervised domain adaptation (UDA), achieving state-of-the-art performance with only 10% labeled target annotations. DAMT-Net employs adversarial multi-task learning to learn domain-invariant and discriminative features [18]. [19] looks at the problem from the lens of pseudo-labeling. NeuroADDA first integrates domain adaptation with active learning in connectomics, surpassing scratch training with a 25-67% reduction in the variation of information [7].

While single source UDA methods have been explored, multi source domain adaption (MDA) approaches remain completely underexplored in connectomics. In natural image segmentation, various MDA methods have been introduced including DCTN [20], MDAN [21], MMN [22], MDDA [23], and MADAN [24]. However, since connectomics datasets vary widely across origin species, imaging protocols, and domain-specific image artifacts [7], existing MDA methods are not directly applicable. In addition, no existing methods combine multi source domain adaptation with active learning. Therefore, this study is the first attempt at combining active learning with multi source domain adaptation by learning ensemble weights over the source domains, training a student model via knowledge distillation, and performing active learning under a constrained annotation budget.

## 3. Methodology

### 3.1. Problem Formulation

Let 𝒳 denote the space of EM images, and let 𝒴 denote the corresponding space of segmentation labels. We are given a target domain 𝒯 ⊂𝒳. In addition, we have access to *m* labeled source domains 𝒮_1_, 𝒮_2_, …, 𝒮_*m*_, where each source domain 𝒮_*i*_ ⊂ 𝒳 ×𝒴 consists of pairs of labeled EM images and their corresponding segmentation masks. Each source domain 𝒮_*i*_ is assumed to originate from a distinct data distribution. For each 𝒮_*i*_, a pretrained segmentation model *F*_*i*_ : 𝒳 → 𝒴 is available, and these models may differ in architecture or representation. The objective is to learn a segmentation function *F* : 𝒳 → 𝒴 that can accurately segment EM images from the target domain 𝒯 by employing information-theoretic principles to transfer knowledge from all available source domains, despite domain shifts and model heterogeneity.

### 3.2. Overview of the MACS Framework

The MACS framework systematically addresses the challenge of segmenting EM images in a target domain by integrating information from multiple heterogeneous source domains. The process begins with multi-source alignment, where MACS employs an information-theoretic strategy based on multi-kernel maximum mean discrepancy (MK-MMD^2^) to optimally combine knowledge from all source domains. This approach learns the relative contribution or weight of each source by minimizing the distributional discrepancy between source and target features, ensuring effective adaptation across diverse data distributions. Once this alignment is established, a unified student model with a fixed architecture is trained. Rather than merging the source models directly, the student model instead learns to reproduce a weighted ensemble of their predictions. This knowledge distillation process uses the learned source weights to combine the outputs from each model, enabling the student to integrate complementary information from all sources in a unified manner. Next, MACS employs active learning to strategically select which unlabeled target images should be annotated next. Active learning in MACS is driven by analytic uncertainty estimation, which is applied directly to the unlabeled images in the target domain. Uncertainty is quantified efficiently in a single forward pass through the student model, providing principled confidence scores for each sample. The most informative, high-uncertainty images are then selected for expert annotation. The student model is subsequently fine-tuned on these newly labeled samples using a regularized objective to preserve previously acquired knowledge. Through this integrated process, MACS enables data-efficient segmentation of the target domain by unifying principled multi-domain adaptation with targeted, uncertainty-driven annotation. Figure 1 visually summarizes the MACS. Next, we discuss our approach in details.

**Figure 1:**
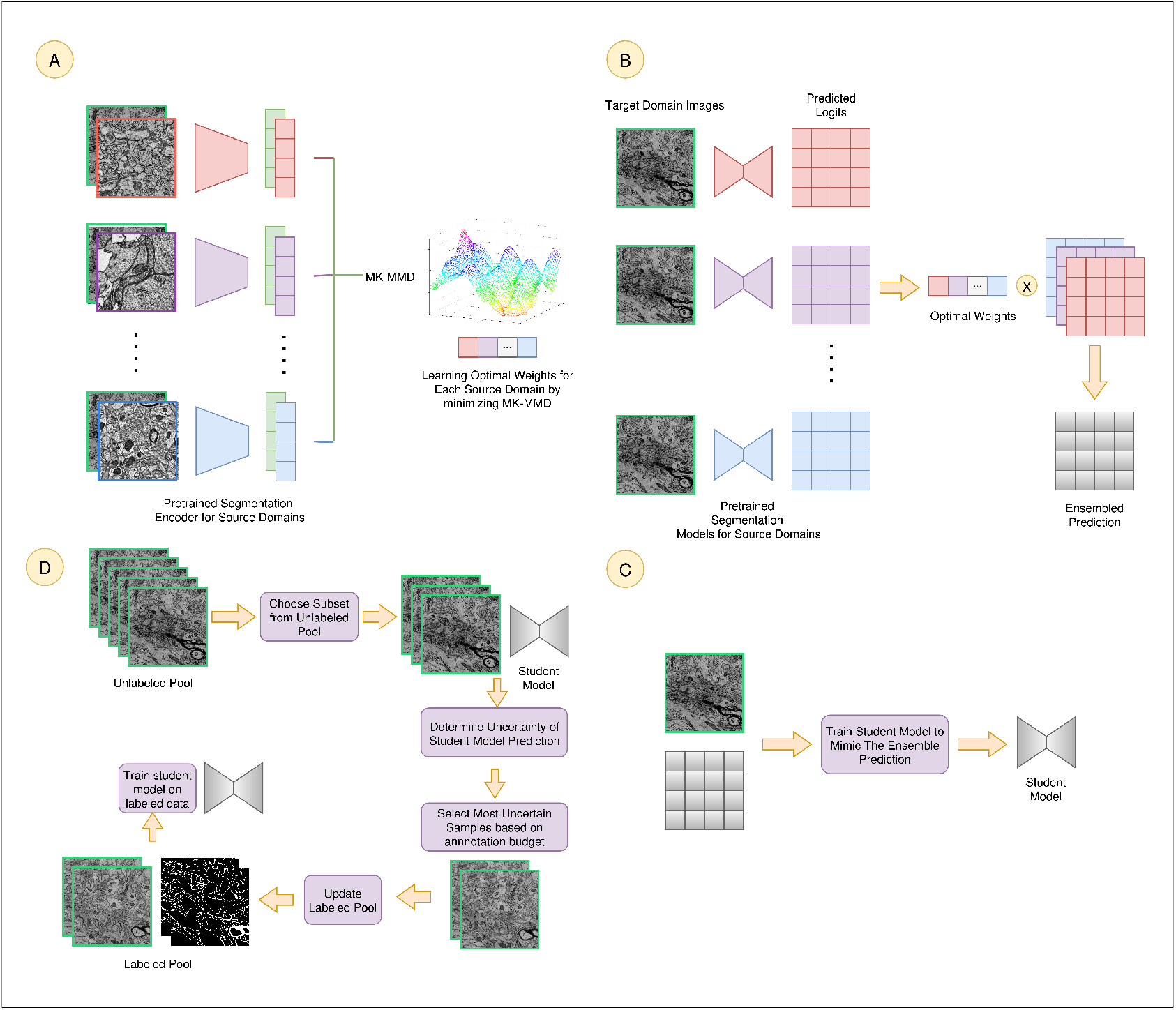
Schematic diagram of MACS.**A.** MACS combines knowledge from multiple heterogeneous source domains by learning optimal source weights using multi-kernel maximum mean discrepancy and aligning features across all domains. **B**. The optimal weights are used to compute a weighted ensemble prediction from all source models. **C**. A unified student model is then trained by knowledge distillation to mimic this ensemble prediction. **D**. Finally, MACS uses active learning with analytic uncertainty estimation to efficiently select the most informative target samples for annotation, enabling the student model to be effectively fine-tuned and adapted to the target domain.

### 3.3 Multi-Source Domain Alignment via Multi-Kernel MMD

The first stage of MACS aims to align the distributions of multiple labeled source domains 𝒮_1_, …, 𝒮_*m*_ with the unlabeled target domain 𝒯 in a principled, information-theoretic manner.

For each source domain 𝒮_*i*_, we extract feature embeddings for every image *x* ∈ 𝒳 by taking the activations from the final down-sampling layer of the corresponding pretrained model *F*_*i*_ and applying global max-pooling to obtain a compact vector representation. We then define a positive-definite kernel *k*_*i*_(*x, x*^*′*^) on these feature vectors, where *x, x*^*′*^ are arbitrary EM images. Each kernel *k*_*i*_ induces its own Reproducing Kernel Hilbert Space (RKHS), uniquely determined by the feature extractor trained on 𝒮_*i*_. Consequently, these embeddings reside in non-comparable RKHSs, making it mathematically invalid to directly add, average, or compare them across domains.

To overcome this incompatibility, MACS constructs a convex combination of the individual kernels,

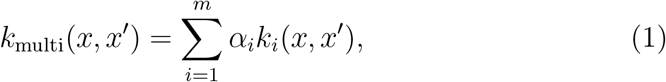

where α_*i*_ *≥* 0 and 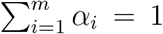. This multi-kernel formulation induces a unified RKHS in which all source and target samples can be meaningfully compared. As a result, the multi-kernel Maximum Mean Discrepancy (MKMMD^2^) can be computed to align the combined source domains with the target domain in a mathematically sound, information-theoretic manner.

The optimal weights ***α*** = [*α*_1_, …, *α*_*m*_] are learned by minimizing the squared multi-kernel Maximum Mean Discrepancy (MK-MMD^2^) between the combined source domains and the target domain:

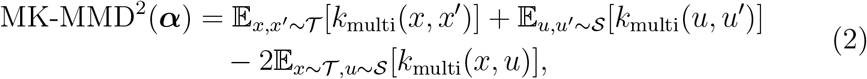

where 𝒮 denotes the union of all source domains. The resulting optimization problem is convex:

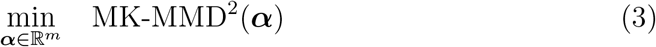

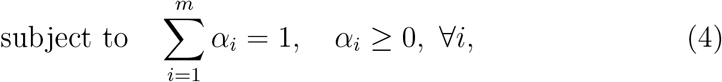

yielding a distribution-weighted kernel that maximally aligns the source and target feature distributions.

### 3.4 Unified Student Model via Knowledge Distillation

Having obtained the optimal kernel weights ***α***^*∗*^, MACS constructs a unified segmentation model *F*_student_ : 𝒳 → 𝒴 suitable for uncertainty estimation and active learning. Direct fusion of source model parameters is infeasible due to architectural heterogeneity. Instead, for each target input *x* ∈ 𝒯, we compute a weighted ensemble of source model predictions:

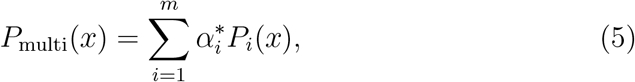

where *P*_*i*_(*x*) = Softmax(*F*_*i*_(*x*)) denotes the probabilistic output of the *i*-th pretrained source model. The student model *F*_student_ is trained to mimic these soft pseudo-labels via knowledge distillation, minimizing the KL divergence loss:

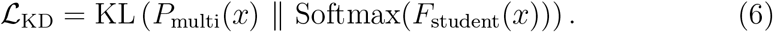

This allows *F*_student_ to consolidate multi-domain expertise into a single architecture that is compatible with downstream adaptation. Hence, MACS is fundamentally agnostic to the architecture of its source models.

### 3.5 Bayesian Uncertainty Estimation via Linearized Laplace Approximation

To provide principled uncertainty quantification suitable for active learning, MACS leverages a Bayesian linearized Laplace approximation focused on the final layer of the student segmentation model. After the student model *F*_student_ is trained, we denote its final-layer weight matrix as *W* ∈ ℝ^*C×d*^, where *C* is the number of segmentation classes and *d* is the dimensionality of the penultimate layer feature space. For any input image *x ∈* 𝒳, the model first computes a feature vector *z*(*x*) *∈* ℝ ^*d*^ via the penultimate layer. The class

#### Algorithm 1

End-to-End Pipeline of MACS.

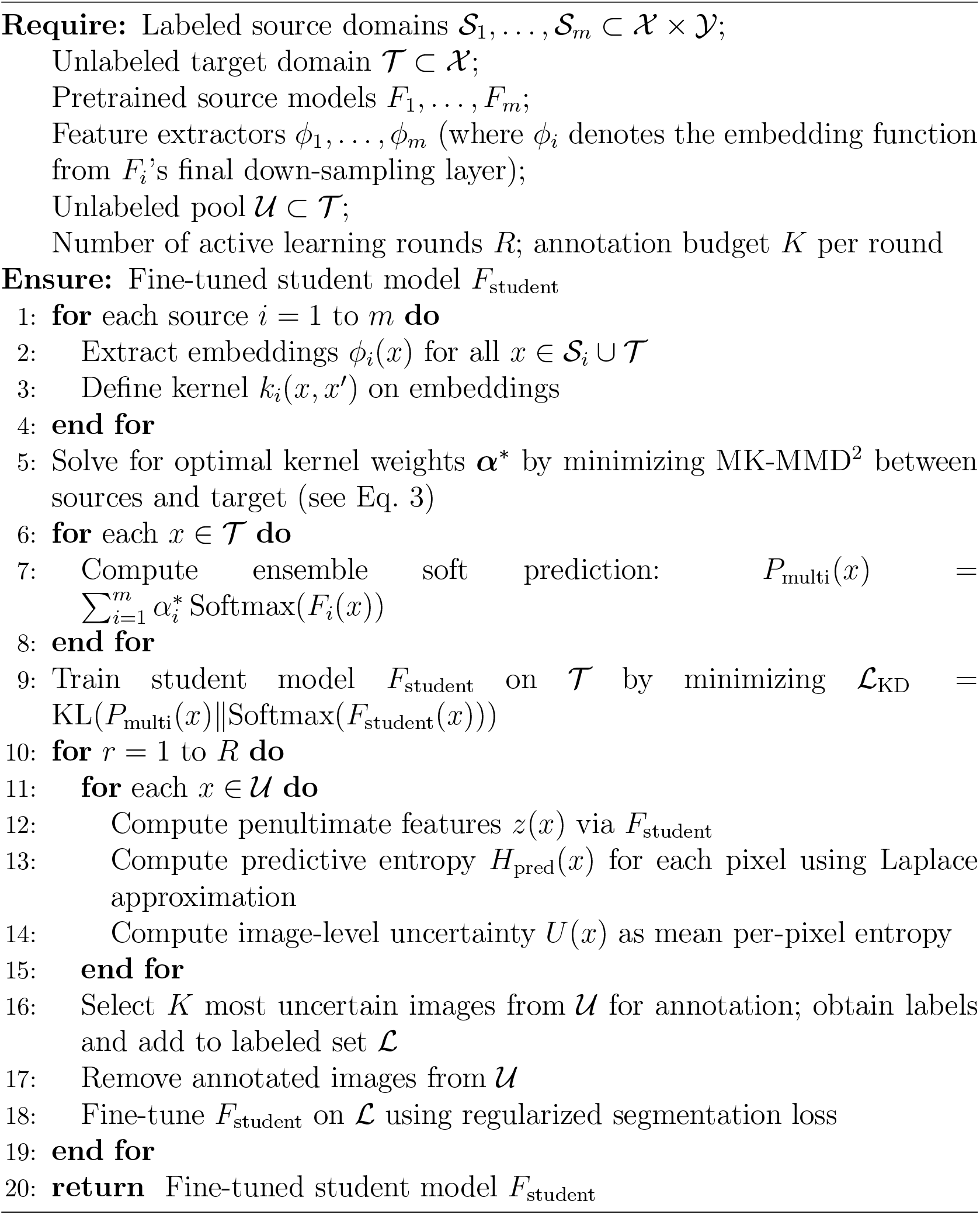

logits are then given by *f* (*x*) = *Wz*(*x*). To capture epistemic uncertainty in a theoretically grounded way, we approximate the posterior distribution over the vectorized final-layer weights as a multivariate Gaussian [25]:

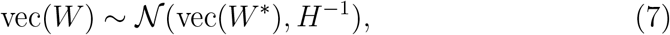

where *W* ^*∗*^ denotes the maximum a posteriori (MAP) estimate of *W*, and *H* is the Hessian of the negative log-likelihood with respect to vec(*W* ), evaluated at *W* ^*∗*^. For computational tractability, we use a diagonal approximation of *H*.

Given this Gaussian posterior over weights and a fixed penultimate feature vector *z*(*x*), the logits for any pixel are also Gaussian-distributed [26]:

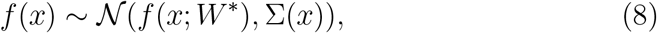

where the mean is *f* (*x*; *W* ^*∗*^) = *W* ^*∗*^*z*(*x*), and the covariance matrix is

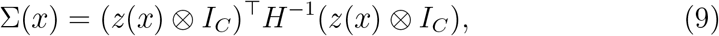

with ⊗ denoting the Kronecker product and *I*_*C*_ the *C*-dimensional identity matrix. This step analytically propagates weight uncertainty to output logits via the penultimate layer’s features, efficiently capturing epistemic uncertainty with no need for Monte Carlo sampling. In contrast to approaches like Monte Carlo Dropout, which require multiple forward passes and substantial computational overhead to estimate uncertainty, MACS yields accurate uncertainty estimates in a single forward and backward pass, making it highly efficient.

For segmentation, we require per-pixel uncertainty estimates. We compute the predictive uncertainty *H*_pred_(*x*) via a closed-form approximation of the expected entropy:

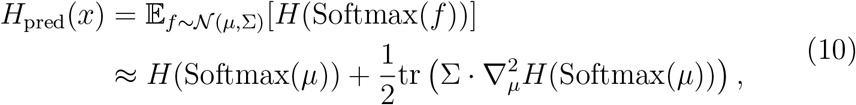

where:

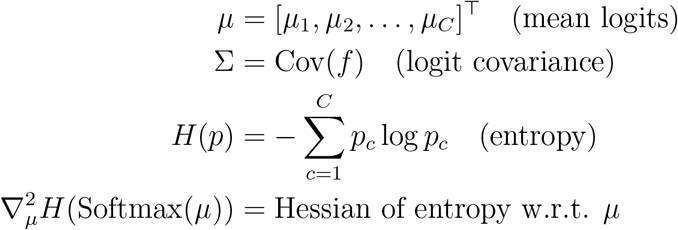

This *H*_pred_(*x*) serves as our uncertainty measure for active learning. Crucially, this closed-form approximation enables efficient uncertainty estimation in just a single forward-backward pass which eliminates the need for costly Monte Carlo dropout while maintaining theoretical rigor.

To obtain an image-level uncertainty score *U* (*x*) for active learning, we aggregate the per-pixel entropy across all *N* pixels in the image:

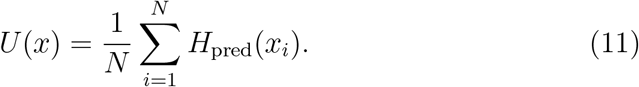

This score reflects the average epistemic uncertainty across the entire segmentation mask, highlighting images for which the model is least confident.

### 3.6 Active Learning and Fine-Tuning

The computed uncertainty scores *U* (*x*) over the unlabeled target pool 𝒰 are then used to drive active learning. Specifically, we select the top-*K* most uncertain images according to *U* (*x*) for expert annotation, thus forming a new labeled set ℒ. The student model *F*_student_ is then fine-tuned on by minimizing a regularized segmentation objective:

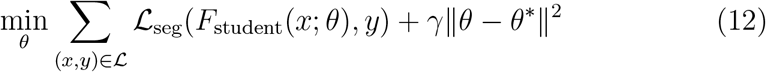

where ℒ _seg_ is the standard segmentation loss that is a combination of binary cross entropy loss, dice loss and focal loss, *θ* denotes the parameters of *F*_student_, *θ*^*∗*^ is the initial parameter set before fine-tuning, and *γ* controls the strength of regularization to preserve previously learned knowledge. Algorithm 1 describes all the steps of our proposed MACS framework.

## 4. Experimental Results

### 4.1 Experimental Setup

We evaluated MACS on nine distinct domains of EM brain tissue datasets: Mouse Somatosensory Cortex (SNEMI3D) [27], Mouse Visual Cortex (MICrONS) [28], Fruit Fly Whole Brain (Flywire) [29], C. elegans Dauer Stage, Mouse Cerebellum (Dhanyasi-P14), Berghia Nerve Cord, Octopus Vertical Lobe Glia Deep Neuropil, Octopus Vertical Lobe SFL Tract, and Whole Mouse Brain [30]. The following Datasets section contains details of these nine datasets. We compare MACS against six representative baselines, organized into three categories:

1. **Domain adaptation:** *NeuroADDA* [7] is a state-of-the-art single-source domain adaptation method.
2. **Connectomics-specific segmentation models:** *PyTorch Connectomics* [31] is a widely-used framework for EM segmentation in connectomics. *FusionNet* [32] is a connectomics-specific convolutional model for EM segmentation. *RETINA* [33] is another baseline pretrained on large-scale EM images and fine-tuned for task-specific segmentation.
3. **Foundation and non-adaptation models:** *Segment Anything Model (SAM)* [34] is a promptable, general-purpose foundation segmentation model. Scratch training refers to supervised training on the target domain alone, without adaptation or external pretraining, isolating the impact of domain adaptation.

We focus on the challenging task of two-dimensional cell membrane segmentation, a critical step in the connectomics pipeline. Our evaluation metrics are the Variation of Information (VI), Dice coefficient, Intersection over Union (IoU), F1 score, and Recall.

All experiments are conducted five times and on a cluster equipped with four NVIDIA RTX A4500 GPUs. For the segmentation backbone, we employ a standard fully convolutional U-Net architecture [35], which is widely used for biomedical image segmentation tasks due to its effectiveness in capturing multi-scale context. For all experiments, we performed active learning over 6 iterations, with 5 training epochs per iteration and a batch size of 4. Each domain is treated in turn as the target, with all remaining domains serving as potential sources for adaptation or transfer experiments. We used the source code provided in the GitHub repository of the baseline models and trained and tested on the exact same data split used for MACS. Since the official implementation of NeuroADDA is not publicly available, we faithfully reimplemented the method based on the details provided in the original paper. For scratch training, we trained the same U-Net architecture using only the target domain training data.

### 4.2. Datasets

We perform a 70:15:15 split of each dataset into training, validation, and test sets, respectively. We employ a patch-based training strategy for all membrane segmentation experiments. Each EM image is divided into overlapping patches of size 256 *×* 256 pixels, with a stride of 128 pixels between adjacent patches. All images and segmentation masks are preprocessed to grayscale, and mask patches are binarized to ensure a consistent foreground-background labeling. Table 1 summarizes the nine datasets used in our study.

**Table 1:** Summary of membrane segmentation datasets used in this study. The Train and Test columns indicate the number of image patches extracted from each dataset split.

| Domain | Species | Brain Region / Tissue | Train | Test |
| --- | --- | --- | --- | --- |
| SNEMI3D | Mouse | Somatosensory Cortex | 3430 | 735 |
| MICrONS | Mouse | Visual Cortex | 42238 | 9114 |
| Flywire | Drosophila | Whole Brain | 35133 | 7595 |
| C. elegans Dauer | C. elegans | Dauer Stage | 5641 | 1025 |
| P14 Cerebellum | Mouse | Cerebellum | 7688 | 3844 |
| Berghia Connective | Berghia | Nerve Cord (Connective) | 436 | 94 |
| Octopus Glia | Octopus | Vertical Lobe Glia Neuropil | 7514 | 2890 |
| Octopus SFL Tract | Octopus | Vertical Lobe SFL Tract | 3072 | 1024 |
| Whole Mouse Brain | Mouse | Whole Brain | 5075 | 1725 |

### 4.3. MACS Achieves Superior Performance Over Connectomics-Specific, Foundation, and Domain Adaptation Baselines

Table 2 presents the mean segmentation performance across nine connectomics domains. For the two domain adaptation methods MACS and NeuroADDA, we report their performance at 100% annotation budget. MACS achieves the highest scores in IoU (0.665), Dice (0.752), and F1 (0.751), outperforming all baselines by large margins. Notably, MACS improves over the strongest adaptation baseline, NeuroADDA, by 34.9% (IoU), 28.3% (Dice), and 28.4% (F1). Compared to the best-performing connectomics-specific model (FusionNet), the improvements are even greater - 8.0% (IoU), 38.7% (Dice), and 38.6% (F1). Furthermore, MACS achieves the lowest variation of information (VI 0.309), representing a 33% reduction relative to the best baseline (NeuroADDA, VI 0.459). For all core metrics, NeuroADDA consistently ranks as the second-best model, followed closely by FusionNet. Although RETINA obtains the highest recall (0.900), this result is due to oversegmentation and comes at the cost of much lower IoU and Dice. We report the performance metrics for each domain in Figure 2.

**Table 2:** Performance Comparison Across All Domains.

| Model | IoU | Dice | F1 | Recall | VI |
| --- | --- | --- | --- | --- | --- |
| FusionNet | 0.421 | 0.542 | 0.542 | 0.579 | 0.462 |
| NeuroADDA | 0.493 | 0.586 | 0.585 | 0.780 | 0.459 |
| PyTC | 0.389 | 0.540 | 0.540 | 0.675 | 0.460 |
| RETINA | 0.136 | 0.229 | 0.229 | **0.900** | 0.636 |
| SAM | 0.193 | 0.311 | 0.311 | 0.859 | 1.236 |
| MACS | **0.665** | **0.752** | **0.751** | 0.813 | **0.309** |

**Figure 2:**
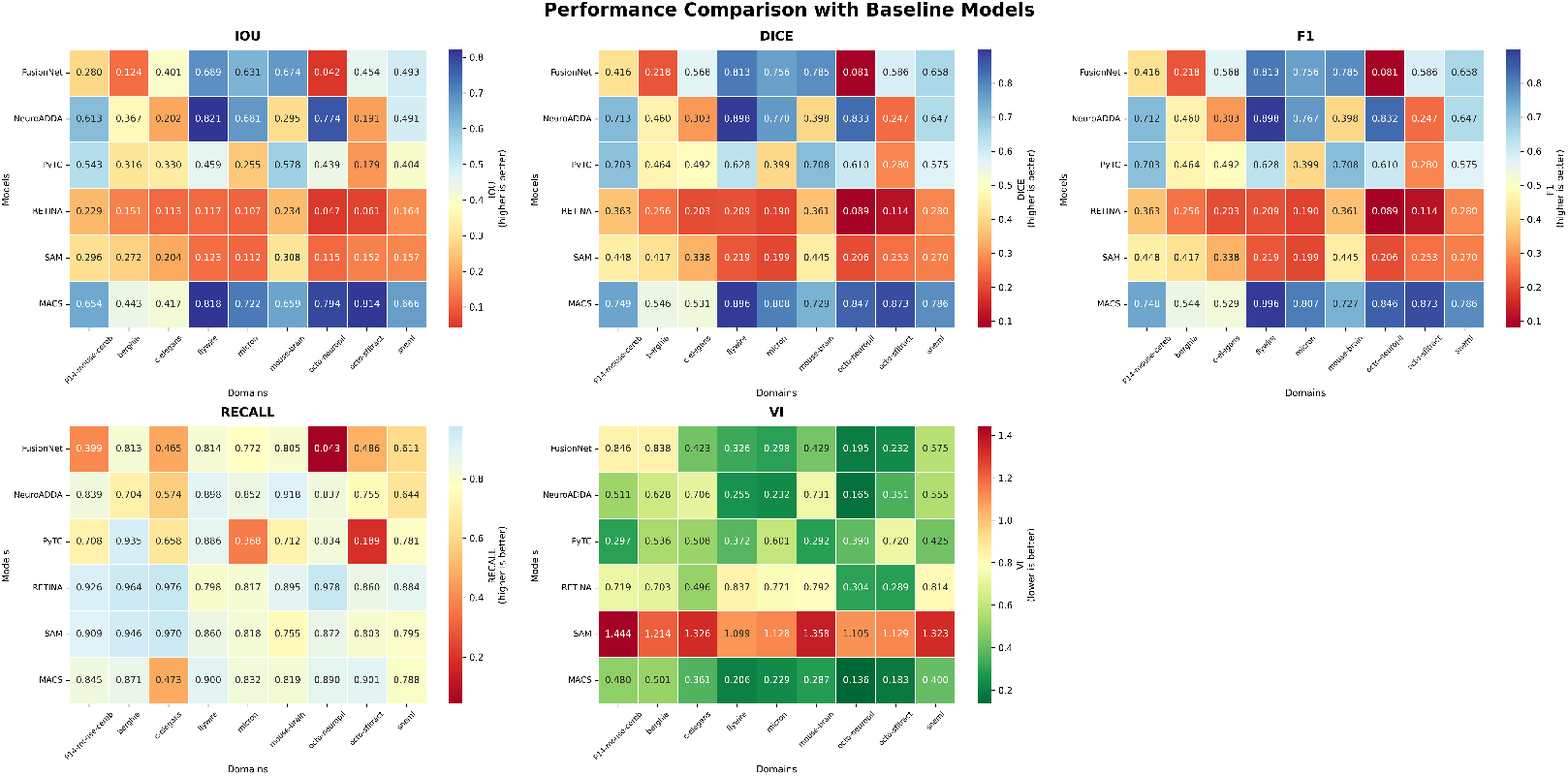
Per-domain and per-metric performance comparison of all models. Each entry reports the score for a given model and metric on an individual domain.

We further evaluate MACS against the state-of-the-art single domain adaptation method NeuroADDA as well as scratch training, across a spectrum of annotation budgets corresponding to ten, twenty, forty, sixty, eighty, and one hundred percent of the available labeled training samples in each dataset. MACS consistently achieves superior performance in terms of the Variation of Information metric, as shown in Figure 3 (See Appendix for performance comparison on other metrics.). Averaged across all annotation budgets and domains, MACS delivers an overall mean percentage improvement of 19.72% relative to NeuroADDA. The mean improvement is 5.89% at the lowest annotation budget (ten percent of training data) and 27.72% at the highest annotation budget (one hundred percent of training data). Analyzing domain-wise mean improvements over all annotation budgets, MACS achieves its highest gains in the octo-sfl domain (41.68%), followed by mouse-brain (35.32%), c-elegans (30.66%), fly (17.41%), octo-glia (17.36%), snemi (10.53%), berghia (10.52%), mouse-cerebellum (9.53%), and micron (4.48%). For the lowest annotation budget, MACS outperforms NeuroADDA in berghia (10.24% improvement), c-elegans (5.32%), fly (11.12%), octo-glia (26.55%), and octo-sfl (15.61%), but underperforms in micron (-1.78%), mouse-brain (-9.58%), mouse-cerebellum (-0.02%), and snemi (-4.46%). At the highest annotation budget, MACS demonstrates dramatic improvements in mouse-brain (60.69%), c-elegans (48.83%), octo-sfl (47.76%), snemi (28.07%), fly (19.42%), berghia (20.20%), octo-glia (17.43%), mouse-cerebellum (6.08%), and micron (1.04%). Across all annotation budgets, MACS wins consistently over NeuroADDA in the domains of berghia, fly, octo-glia, and octo-sfl. The best individual improvement is observed in mouse-brain at one hundred percent annotation budget, with a 60.69% increase. The worst individual improvement occurs in mouse-brain at ten percent annotation budget, with a -9.58% decrease. In every experiment, training from scratch consistently yields lower segmentation performance than both MACS and NeuroADDA across all annotation regimes and domains. The reported results are statistically significant, assessed using a paired twosided t-test (*p <* 0.01). Overall, these results highlight the effectiveness of MACS, which sets a new state-of-the-art across all metrics and achieves substantial improvements over both connectomics-specific, adaptation-based, and foundation segmentation models.

**Figure 3:**
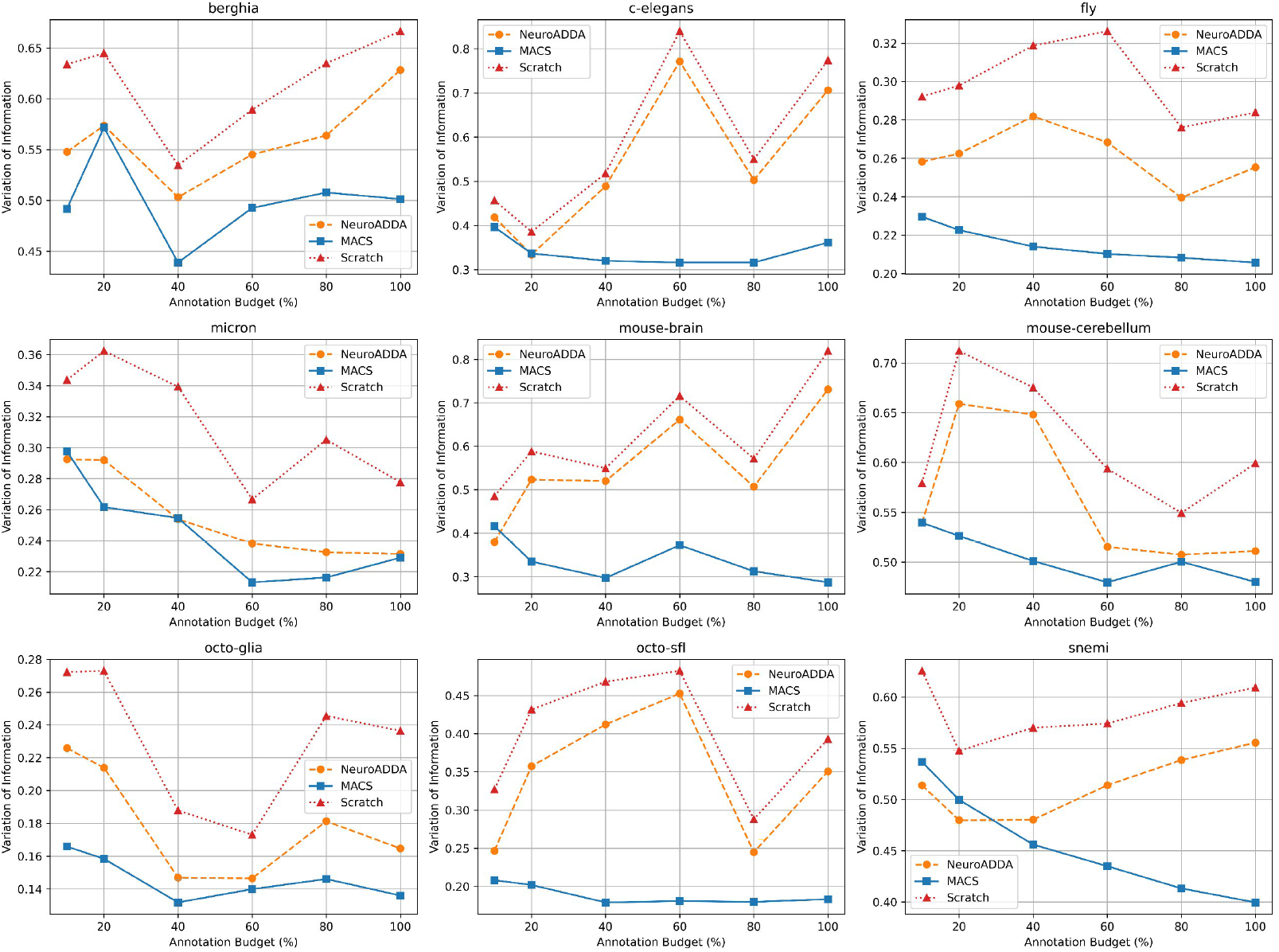
Comparison of segmentation performance (VI) across annotation budgets for MACS, NeuroADDA, and Scratch Training on all domains. MACS consistently outperforms both baselines at all annotation levels.

### 4.4 MACS Supports Heterogeneous Model Architectures Across Different Source Domains

Traditional domain adaptation frameworks in connectomics typically require all source domain models to share an identical architecture [7]. This restriction severely limits practical applicability, since public pretrained models for various domains are rarely standardized, and different datasets may demand distinct model designs for optimal performance. By contrast, MACS is fundamentally agnostic to the architecture of its source models, as described previously. This allows MACS to flexibly incorporate the most effective architecture for each source domain, maximizing transferability and real-world usability. To empirically validate this capability, we conducted experiments where source domains contributed models from three popular families: UNet [35], ResUNet [36], and DeepLab [37]. We report the average Variation of Information (VI) for MACS on each domain, using the best-suited architecture per domain, as shown in Table 3.

**Table 3:** VI scores for MACS using three model architectures (DeepLab, ResUNet, UNet) across nine connectomics domains.

| DeepLab |  | ResUNet |  | UNet |  |
| --- | --- | --- | --- | --- | --- |
| Domain | VI | Domain | VI | Domain | VI |
| berghia | 0.3250 | c-elegans | 0.2100 | fly | 0.2151 |
| octo-glia | 0.1172 | mouse-brain | 0.2211 | micron | 0.2454 |
| octo-sfl | 0.1500 | snemi | 0.3220 | mouse-cerebellum | 0.5045 |

### 4.5. MACS Offers Efficient and Scalable Uncertainty Computation With Laplace Approximation

A key advantage of MACS is its support for fast and scalable uncertainty estimation using analytic Laplace approximation, in contrast to traditional Monte Carlo Dropout (MC Dropout) approaches which require repeated stochastic forward passes through the network. The Laplace method yields high-quality uncertainty estimates in a single analytic computation which enables efficient active learning and sample selection. Table 4 summarizes the average computation time for uncertainty scores using Laplace approximation in MACS, compared to MC Dropout with varying numbers of stochastic passes as used in NeuroADDA [7]. The results demonstrate that the Laplace approach is substantially faster and more scalable, being over 4*×* to 24*×* faster than MC Dropout for typical settings.

**Table 4:** Average time (in seconds) required to compute uncertainty scores using Laplace approximation (MACS) and MC Dropout for different numbers of forward passes.

| Method | Setting | Computation Time (s) |
| --- | --- | --- |
| MC Dropout | 10 Passes | 194.09 |
| MC Dropout | 20 Passes | 501.45 |
| MC Dropout | 50 Passes | 1121.64 |
| Laplace Approx. (MACS) | Analytic (Single Pass) | **46.49** |

### 4.6. MACS Learns to Upweight the Most Transferable Source Domains: A LEEP-Based Analysis

To rigorously assess whether MACS assigns higher weights to source domains that are genuinely more transferable to the target, we conduct a systematic analysis using the Log Expected Empirical Prediction (LEEP) score [38], a principled metric for quantifying cross-domain adaptability. For each source-target domain pair, we begin by extracting penultimate-layer features from the target domain using the encoder trained solely on the source. We then train a linear classifier on the source features and labels, and use this classifier to compute, for each target sample *x*_*i*_, the predicted probability *P* (*z* | *x*_*i*_) of belonging to each source class *z*. To quantify the empirical relationship between source and target label spaces, we estimate *P* (*y* |*z*) as the probability that a sample assigned to source class *z* actually corresponds to target label *y* in the target domain, computed as :

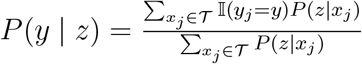

which intuitively reflects how well the where *y*_*j*_ is the ground-truth label of *x*_*j*_ in the target domain. The LEEP score for each source–target pair is then given by : 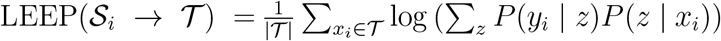 source-trained features align with the target label distribution. A higher LEEP score indicates greater compatibility between the source’s learned representations and the target domain’s labeling structure. Across all targets, we find that the weights 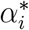 learned by MACS exhibit a strong Spearman rank correlation (0.91) with the corresponding LEEP scores. This analysis demonstrates that MACS consistently upweights sources with high empirical transferability while downweighting less relevant sources, thus providing mechanistic evidence that its information-theoretic weighting strategy selects the most transferable domains in practice.

### 4.7 Case Study

We conducted a qualitative case study on the Flywire as target domain using models trained with 100% annotation budget. Test samples were grouped into easy, moderate, and hard categories based on prediction difficulty. As shown in Fig. 4, MACS consistently produces more contiguous and very less porous membrane segmentations across all difficulty levels. In contrast, single domain adaptation based NeuroADDA often yields fragmented or porous boundaries in moderate and hard cases. These results highlight MACS’s effectiveness in challenging settings.

**Figure 4:**
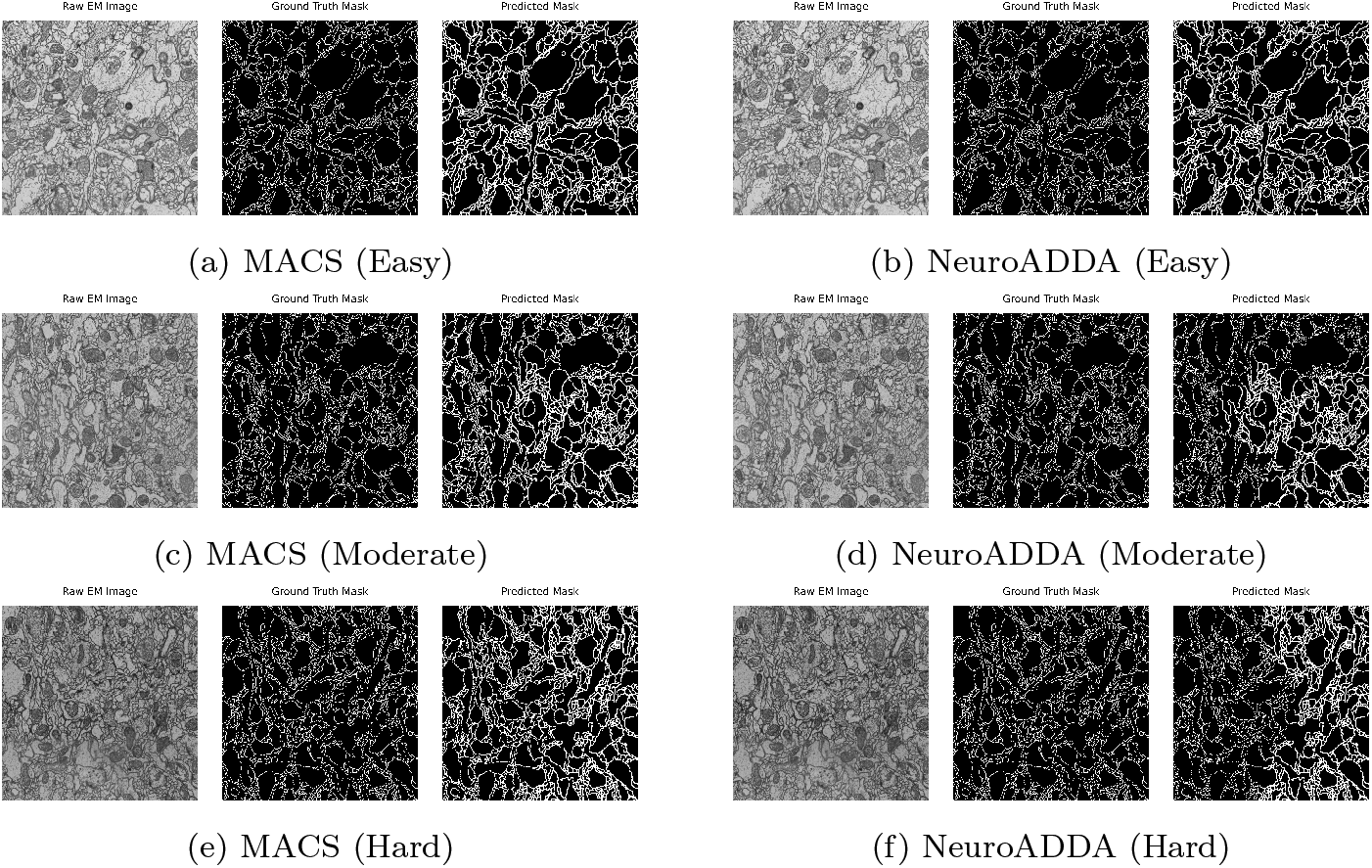
Qualitative segmentation comparison between MACS and NeuroADDA across difficulty levels (easy, moderate, and hard). Each cell shows, from left to right: raw image, ground truth, and predicted masks for the corresponding model and difficulty. Figure size is increased for visual clarity and does not exceed page bounds.

### 4.8. Ablation Study

#### 4.8.1. Impact of the Number of Source Domains on Adaptation Performance

We analyze the impact of increasing source domain diversity in multidomain adaptation. As the number of source domains increases from two to nine, the average VI on the target domain steadily decreases from 0.7100 (two sources) to 0.3262 (all nine), as shown in Figure 5a. This trend demonstrates that leveraging more diverse sources consistently improves adaptation performance, underscoring the value of multi-domain training in MACS.

**Figure 5:**
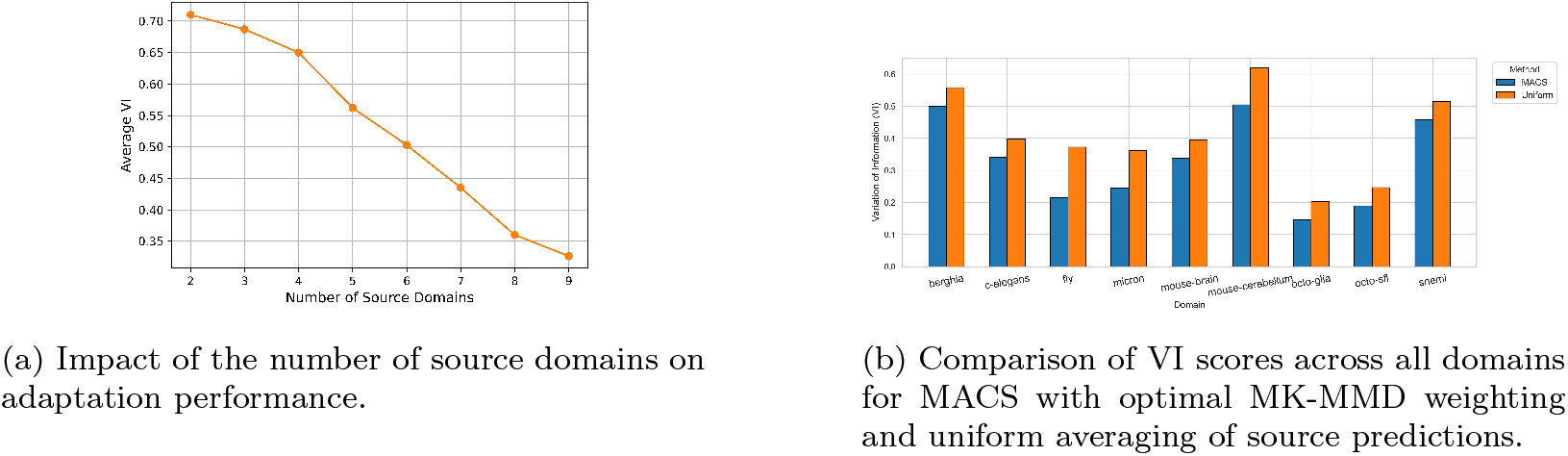
(a) Increasing the number of source domains improves segmentation results. (b) MACS outperforms uniform source averaging across all domains.

#### 4.8.2. Effect of Source Domain Weighting Versus Uniform Averaging

We compare the adaptive MK-MMD based source domain weighting in MACS to uniform averaging of source predictions across all domains. As shown in Figure 5b, MACS consistently achieves lower VI scores than the uniform baseline, with the largest gains for domains such as micron and fly. This experiment demonstrates the benefit of adaptive weighting in multidomain adaptation and further motivates the design of MACS.

## 6. Conclusion

In this work, we present MACS, a novel multi-domain adaptation framework for segmentation in connectomics. Unlike traditional domain adaptation approaches that rely on a single source or enforce architectural uniformity where all source models have the same architecture, MACS is designed to aggregate information from multiple, heterogeneous source domains and model architectures. This flexibility is a direct response to the practical realities of large-scale biomedical imaging, where public models and datasets are diverse, and domain shift is often pronounced.

A central contribution of MACS is its data-driven information theoretic source domain weighting mechanism. By learning interpretable weights over multiple source domains through a convex combination of kernels, MACS is able to prioritize sources that empirically transfer best to the target, as validated by their alignment with LEEP-based transferability scores in our experiment. This approach not only improves segmentation accuracy, but also brings a layer of mechanistic interpretability to domain adaptation, an aspect often lacking in deep learning-based transfer frameworks.

MACS further advances the field by integrating analytic Bayesian Laplace-based uncertainty quantification within its active learning cycle. As supported by our experiment, unlike traditional Monte Carlo dropout, Laplace approximation provides a scalable and computationally efficient means of estimating the uncertainty. This enables more informed sample selection for annotation and ensures efficient learning.

Our extensive empirical study, spanning nine diverse connectomics domains, demonstrates that MACS outperforms both single-domain adaptation method and state-of-the-art connectomics baselines across all key segmentation metrics and annotation regimes. The framework’s architecture-agnostic design also facilitates seamless integration of future deep learning advances, making MACS highly adaptable as new models and domains emerge.

Despite its strengths, a current limitation of MACS is its dependence on the overlap between source and target domain structures – performance may degrade when the target contains novel morphologies absent from all sources. Addressing this, future work will focus on equipping MACS with mechanisms for open-set adaptation, domain generalization, and self-supervised representation learning to enable robust segmentation even under extreme domain shift or unseen biological structures. Further, we plan to extend MACS by incorporating semi-supervised and contrastive learning objectives, and by evaluating generalization to new imaging modalities.

In summary, MACS represents a significant step forward in the integration of modern machine learning methodology into large-scale connectomics segmentation. By unifying multi-domain adaptation, efficient uncertainty quantification, and model-agnostic transfer, MACS lays a foundation for robust, scalable, and interpretable ML approaches in neuroscience and beyond.

## Appendix A. Hyperparameter Details of MACS

The hyperparameter details of MACS are presented in Table A.5.

## Appendix B. Evaluation Metrics

To assess the performance of the EM image segmentation models, we utilize a comprehensive set of metrics—Dice Score, Intersection over Union (IoU), Precision, Recall, and Variation of Information (VI). These metrics evaluate both pixel-wise accuracy and structural consistency, providing a robust framework for evaluating segmentation quality.

### Appendix B.1. Dice Score

The Dice Score measures the overlap between the predicted segmentation *P* and the ground truth *G*:

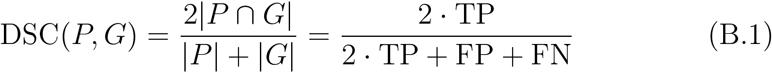

Here, TP, FP, and FN denote true positives, false positives, and false negatives, respectively. Dice Score ranges from 0 (no overlap) to 1 (perfect overlap). Since, we are evaluating on a binary segmentation task, Dice Score is also equivalent to F1-score.

### Appendix B.2. Intersection over Union (IoU)

IoU, or Jaccard Index, evaluates the overlap relative to the union of predicted and ground-truth segmentations, particularly sensitive to boundary accuracy:

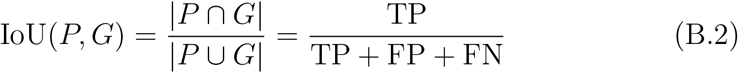

IoU ranges from 0 to 1, with higher values indicating better segmentation.

### Appendix B.3. Recall

Recall measures the proportion of ground-truth pixels correctly identified, ensuring structures are completely captured:

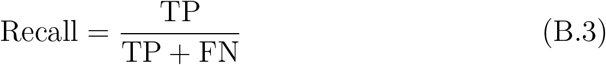

A high Recall score (near 1) reflects minimal under-segmentation, critical for complete reconstruction.

**Table A.5:** Summary of Hyperparameters for MACS Framework.

| Hyperparameter | Value/Default | Description |
| --- | --- | --- |
| *Data / Training Control* |  |  |
| data_root | data | Root directory for datasets |
| source_domains | - | List of source domain names |
| source_ckpts | - | List of paths to pretrained source model checkpoints |
| target_domain | - | Target dataset/folder name |
| out_path | - | Path to save the final trained model |
| batch_size | 8 | Batch size for all DataLoader instances |
| active_iters | 10 | Number of active learning iterations |
| annot_budget | 100 | Annotation budget as percent of target pool. Varies depending on experiment. |
| train_epochs_per_iter | 5 | Number of epochs for fine-tuning on labeled data per iteration |
| device | cuda/cpu | Computation device selection |
| patch_size | 256 | Patch size for image and mask extraction |
| stride | 128 | Stride for sliding window patch extraction |
| num_workers | 2 | Number of worker processes for DataLoader |
| *Model and KD Training* |  |  |
| in_ch | 1 | Number of input channels for UNet |
| out_ch | 1 | Number of output channels for UNet |
| lr | $1 \times 10^{-4}$ | Learning rate for Adam optimizer |
| epochs (KD) | 5 | Number of epochs for KD student training |
| features | [64, 128, 256, 512] | List of feature map sizes for each encoder/decoder stage |
| kernel_size | 3 (conv), 2 (pool/up) | Convolution kernel size (3), max pooling and upsample kernel size (2) |
| activation | ReLU | Activation function after each conv layer |
| normalization | BatchNorm2d | Type of normalization after each convolution |
| final_conv | kernel size 1 | Final output convolution for segmentation mask |
| skip connections | Yes | Skip connections (concatenation in decoder) |
| *Loss Functions* |  |  |
| Dice loss smooth | 1.0 | Smoothing constant for Dice loss |
| Focal loss $\alpha$ | 0.8 | Focal loss weighting parameter |
| Focal loss $\gamma$ | 2 | Focal loss focusing parameter |
| BCEWithLogitsLoss | - | Used as-is (no custom hyperparams) |
| *MK-MMD/Active Learning* |  |  |
| kernel (MK-MMD) | rbf | Kernel function for MK-MMD (default is RBF) |
| lambda_entropy | 0.01 | Entropy regularization coefficient for MK-MMD ($\lambda_{entropy}$) |
| K | derived | Number of samples acquired per active learning round |
| *Laplace Uncertainty* |  |  |
| Small epsilon (uncertainty) | 1e-6 | Epsilon for numerical stability in uncertainty/soft labels |

### Appendix B.4. Variation of Information (VI)

VI, an information-theoretic metric, measures the distance between predicted and ground-truth segmentations by comparing pixel clustering, ideal for complex structures:

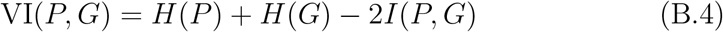

Here, *H*(*P* ) and *H*(*G*) are the entropies of *P* and *G*, and *I*(*P, G*) is their mutual information.

Lower VI values indicate greater similarity between segmentations.

## Appendix C. Comparison of Segmentation Performance between MACS and Single Domain Adaptation in terms of Dice Score, IoU, and Recall

In the main paper, we compared segmentation performance between MACS and single-domain adaptation using Variation of Information (VI). Here, we extend this comparison to three additional metrics: Dice Score, Intersection over Union (IoU), and Recall. The results for each metric are shown in Figure C.6, Figure C7, and Figure C.8, respectively. Across all metrics and annotation budgets, MACS consistently outperforms single-domain adaptation (NeuroADDA), while training from scratch results in the lowest performance.

**Figure C6:**
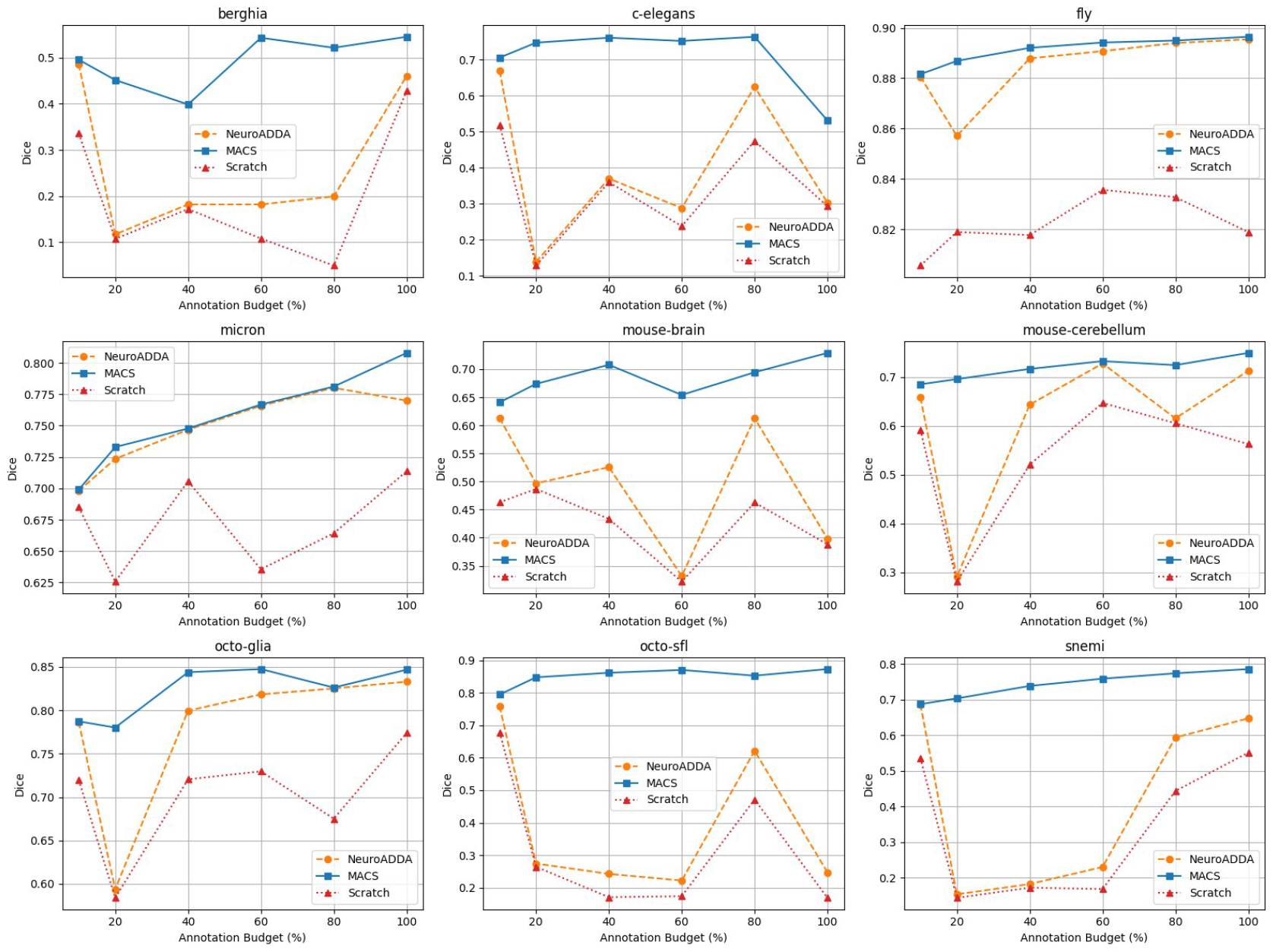
Comparison of segmentation performance (Dice score) across annotation budgets for MACS, NeuroADDA, and Scratch Training on all domains.

**Figure C7:**
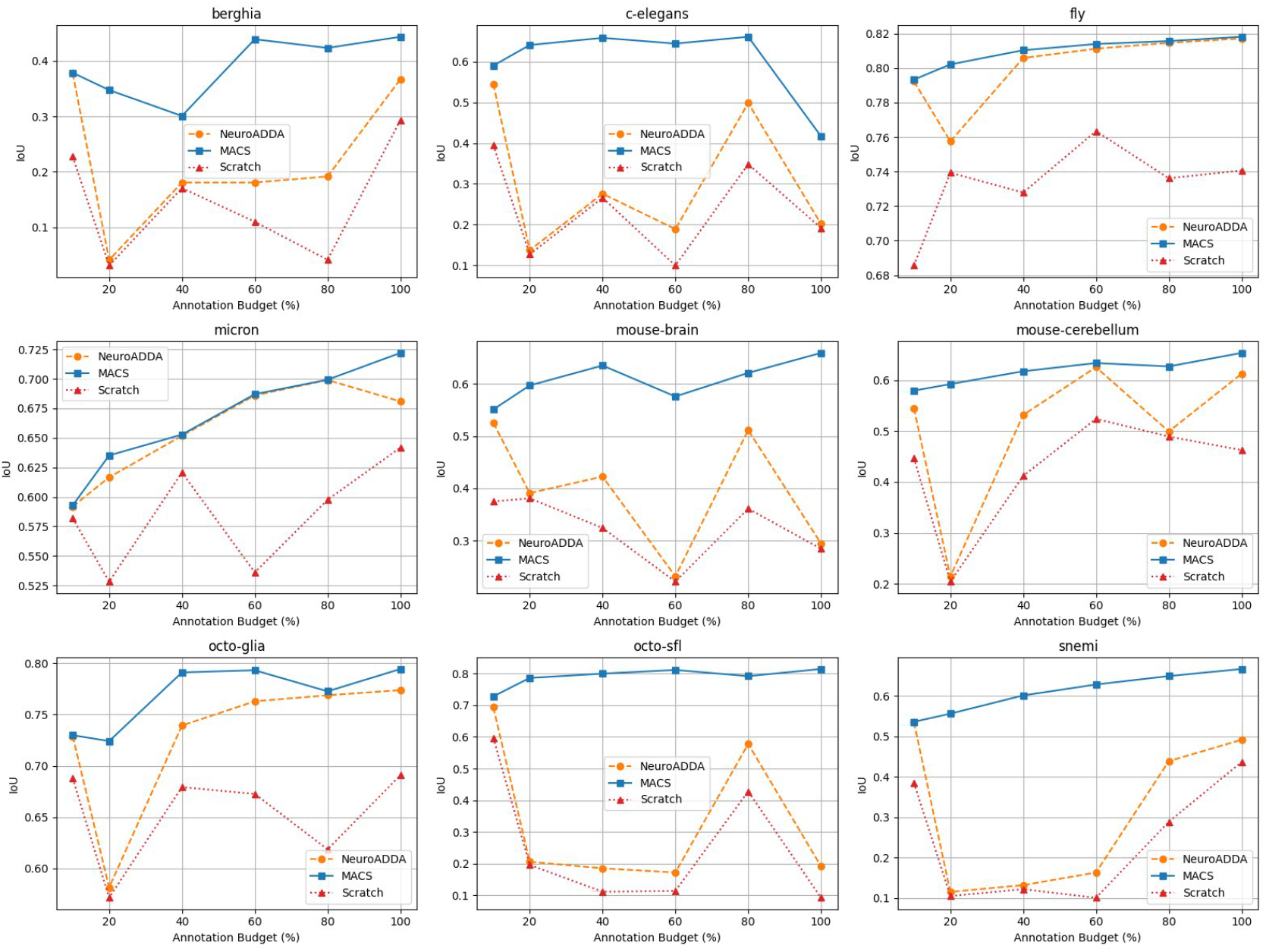
Comparison of segmentation performance (IoU) across annotation budgets for MACS, NeuroADDA, and Scratch Training on all domains.

**Figure C8:**
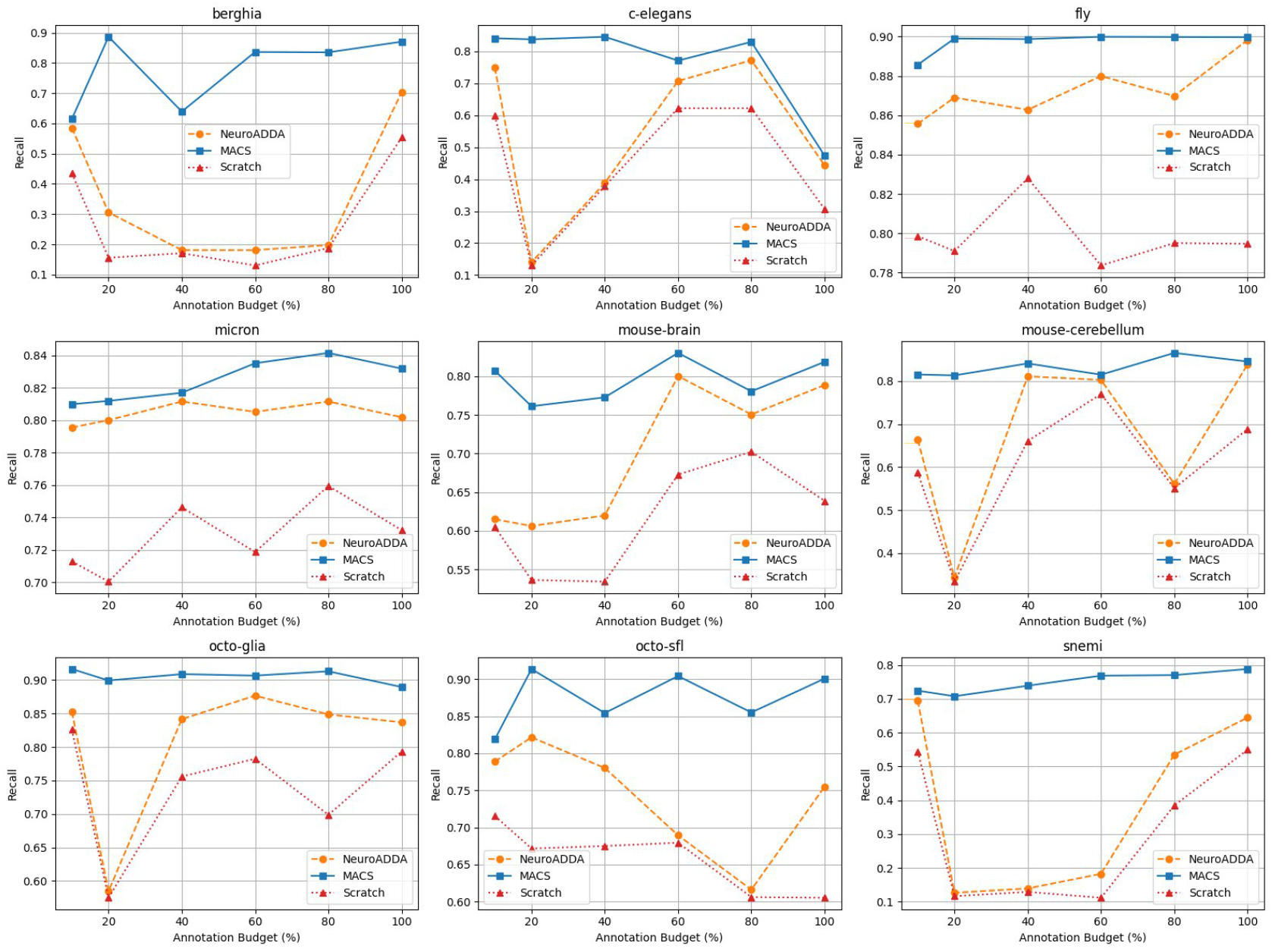
Comparison of segmentation performance (Recall) across annotation budgets for MACS, NeuroADDA, and Scratch Training on all domains.

